# Non-Invasive Photoacoustic Imaging of Cerebral Oxygenation and Hemoglobin Content in Awake Mice

**DOI:** 10.1101/2025.10.02.679935

**Authors:** Juri Aparicio Arias, Chrystel Lafont, Philippe Trochet, Dieter Fuchs, Pierre Sicard

**Affiliations:** IPAM, Biocampus, INSERM/CNRS/Université de Montpellier, Montpellier France; INSERM/CNRS/Université de Montpellier, IGF, Montpellier, France; FUJIFILM VisualSonics Inc., 1114 AB Amsterdam, The Netherland; INSERM/CNRS/Université de Montpellier, PhyMedExp, IPAM/Biocampus, Montpellier, France

**Keywords:** Awake cerebral activity, PA imaging

## Abstract

**Introduction:** Investigating cerebral oxygen saturation dynamics in awake animal models remains technically challenging due to motion artifacts and anesthesia-related biases. Here, we introduce a novel high-resolution ultrasound-photoacoustic (PA) imaging approach enabling real-time, non-invasive monitoring of deep cerebrovascular oxygenation dynamics in awake mice with intact skulls.

**Materials and Methods:** Swiss male and female mice (n = 5–6) were head-fixed using a customized holder adapted to the Neurotar Mobile HomeCage floating platform. High-resolution ultrasound combined with PA imaging (VevoLAZR-X, VisualSonics) was used to discriminate oxyhemoglobin, deoxyhemoglobin, and total hemoglobin in multiple brain regions. Cerebrovascular responses were assessed under three paradigms: (i) baseline awake state vs. 2% isoflurane anesthesia, and (ii) right whisker stimulation to probe sensory-driven hemodynamics.

**Results:** PA imaging successfully resolved deep-brain oxygenation in awake, intact-skull mice. Under isoflurane anesthesia, we observed a rapid and transient increase in cerebrovascular sO□ (p < 0.01). During whisker stimulation, we detected robust, region-specific increases in total hemoglobin, reflecting localized neurovascular coupling in awake mice.

**Conclusions:** This study establishes high-resolution PA imaging as a powerful, non-invasive tool to monitor cerebrovascular oxygenation dynamics in awake mice. By integrating baseline, anesthetic, and sensory paradigms, we demonstrate its potential to dissect neurovascular physiology without the confounding effects of anesthesia. These findings provide new opportunities for preclinical neuroscience research and translational applications investigating cerebral oxygen metabolism.

## INTRODUCTION

Accurate assessment of cerebral oxygenation dynamics is essential for understanding brain physiology and its alterations in neurological disorders. Traditional approaches for monitoring cerebrovascular oxygenation, such as functional ultrasound (fUS) (1), blood-oxygen-level-dependent (BOLD) functional magnetic resonance imaging (fMRI) (2), and intravital imaging microscopy (3), have significantly advanced our ability to probe neurovascular function. However, each modality faces important limitations: BOLD fMRI requires costly infrastructure, long acquisition times, and is highly sensitive to motion artifacts; fUS involves relatively complex acquisition and processing pipelines; and most intravital microscopy techniques are restricted by shallow imaging depths or require invasive cranial windows, limiting their use for longitudinal studies in awake animals.

Photoacoustic (PA) imaging has emerged as a powerful alternative for functional brain studies, combining the molecular sensitivity of optical absorption with the penetration depth and spatial resolution of ultrasound (4,5). This hybrid technique enables label-free, real-time quantification of oxyhemoglobin, deoxyhemoglobin, and total hemoglobin in vivo, providing a direct readout of cerebrovascular oxygenation. Importantly, PA imaging can be performed non-invasively through the intact skull, making it particularly suitable for longitudinal and translational studies. It has already shown promise in detecting brain hypoxia after traumatic brain injury (6) and in tracking oxygenation trajectories during ischemia–reperfusion following cardiac arrest and resuscitation (7). Despite these advantages, the application of high-resolution PA imaging in awake, behaving small-animal models has remained technically challenging, primarily due to the lack of stabilization strategies compatible with naturalistic behavioral contexts. To overcome these limitations, we implemented a high-resolution ultrasound-photoacoustic imaging platform specifically designed for awake mice. By integrating a customized head-fixation system with the Neurotar Mobile HomeCage environment, we achieved stable and minimally stressful acquisition conditions, enabling the investigation of cerebrovascular dynamics without anesthesia or invasive skull modifications. This experimental paradigm allows systematic characterization of cerebrovascular function under three complementary conditions: (i) baseline awake states, (ii) modulation by isoflurane anesthesia, and (iii) sensory-driven hemodynamic responses elicited by whisker stimulation.

In this work, we present a comprehensive characterization of cerebrovascular oxygenation dynamics in awake mice using this non-invasive, high-resolution PA imaging approach.

## Materials and Methods Animals

Adult Swiss male and female mice (n = 5–6, aged 8–12 weeks, weighing 25–28 g) were used in this study. Animals were housed under standard conditions with a 12 h light/dark cycle, controlled temperature (22 ± 1 °C), and ad libitum access to food and water. All experimental procedures were conducted in accordance with the European Directive 2010/63/EU and approved by the local Institutional Animal Care and Use Committee under the number #45060-2023100417365343 v4. The animal habitat was held at a constant 22 ± 1ºC with 58 ± 1% humidity. Animals had access to food and water at ad libitum. (8)

### Awake Imaging Setup

To enable stable, high-resolution imaging in awake conditions, mice were head-fixed using a custom-designed holder mounted on a Neurotar Mobile HomeCage® platform (Neurotar, Helsinki, Finland). The Mobile HomeCage is a floating platform that allows free locomotion while minimizing head motion, providing a naturalistic behavioral context during imaging. The customized head-fixation apparatus ensured reproducible alignment of the imaging plane across sessions and animals (Figure 1 A). Cranial window is implanted according to published protocols (9) (Figure 1B). Prior to imaging, mice were habituated to the setup for 15 consecutive days (10–15 min per session) to reduce stress and minimize motion artifacts during acquisitions (Figure 1C-E).

**Figure 1.**
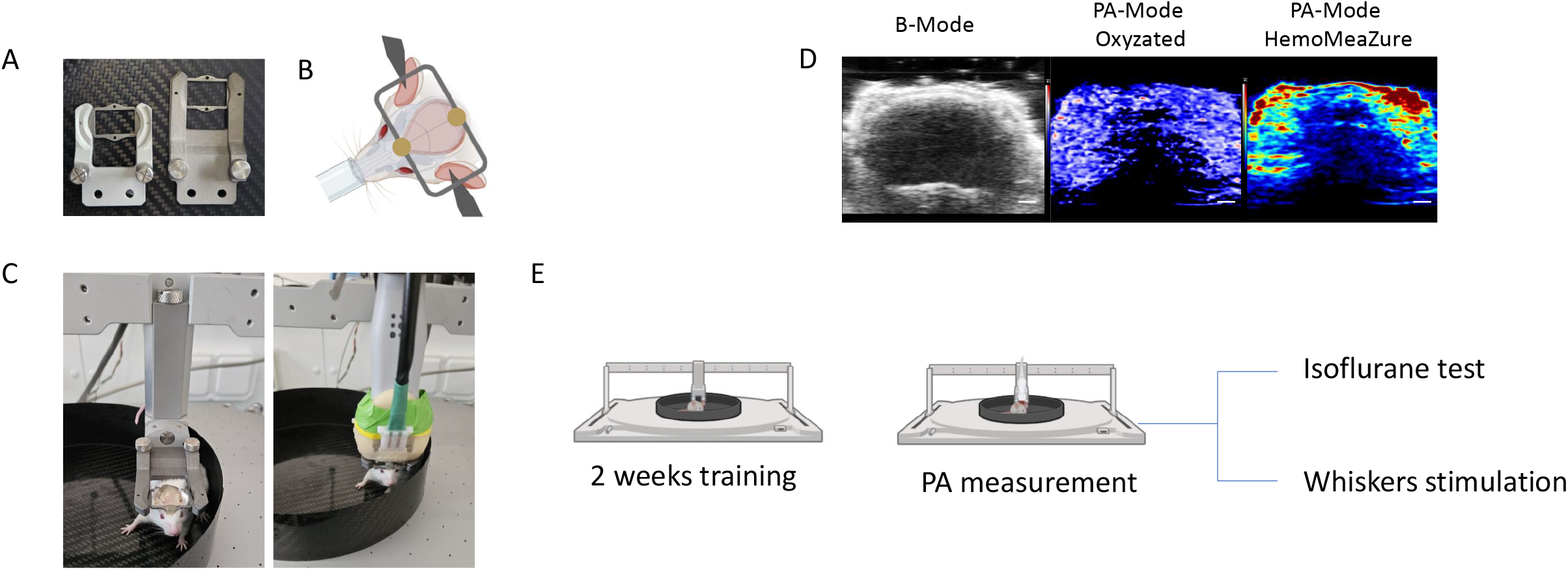
Neurotar Mobile HomeCage and photoacoustic imaging setup for awake mouse brain imaging. **Panel A** Customized head-plate holder designed for photoacoustic imaging (right) compared to the conventional holder used for functional ultrasound (fUS) imaging. **Panel B** Surgical procedure for affixing head plates (model 14) onto the intact skull of mice under isoflurane anesthesia. **Panel C** Awake mouse positioned in the Neurotar Mobile HomeCage with or without the combined PA-US probe. **Panel D** Representative high-resolution ultrasound images and corresponding photoacoustic reconstructions enabling discrimination between oxyhemoglobin and deoxyhemoglobin (PA-mode OxyZated) and visualization of total hemoglobin (PA-mode HemoMeaZure). **Panel E** Schematic representation of the experimental protocol.

### Ultrasound and Photoacoustic Imaging

As previoulsly described (10) PAI was performed with a LAZR-X (Nd:YAG 680–2000□nm) concurrently with the Vevo 3100 (Vevo® LAZR-X, FUJIFILM VisualSonics Inc.) using a 15–30□MHz MX250D probe to assess brain tissue oxygenation. Briefly, mice were anesthetized with 2 % isoflurane (1.0 L/min) and kept at 37 °C. Hair was removed with depilatory cream, and bubble-free ultrasound gel was applied. PAI was acquired at 750/850 nm, with B-mode (US) using standard settings (PA gain 40 dB, depth 22 mm, width 21 mm). Each scan (∼4–5 min) yielded ∼300 frames, which analyzed offline in using the Vevo LAB software (v5.9.1) (FUJIFILM VisualSonics Inc.) for sO□ and HbT.

### Cerebrovascular responses protocol

Baseline awake state – Spontaneous cerebral oxygenation was recorded in awake, head-fixed mice at rest to establish reference hemodynamic levels. To evaluate the effects of anesthesia on cerebral oxygenation, awake mice were exposed to 2% isoflurane in medical air delivered via a nose cone. PA acquisitions were performed continuously during transitions between the awake state and isoflurane exposure to capture transient oxygenation dynamics.

To probe neurovascular coupling, the right whiskers were stimulated using a cotton swab for 20-30 seconds while PA imaging was performed over the brain. The timing of stimulus delivery was synchronized with image acquisition to resolve stimulus-evoked hemodynamic changes.

### Data Processing and Quantification

PA signals were reconstructed and spectrally unmixed using Vevo LAB software (VisualSonics) to calculate regional cerebral oxygen saturation (sO□) and Total hemoglogin (HbT). Region-of-interest (ROI) analyses were performed in predefined cortical and subcortical areas to extract dynamic oxygenation profiles from the left and right hemisphere.

### Statistical analysis

All statistical analysis was performed in prism 8.1.1 (GraphPad Software) and the data are presented as mean ± standard deviation (SD). For comparisons between groups with repeat measurement comparisons, we checked for normality with a Shapiro–Wilk test. Normally distributed data were compared with two-tailed paired Student’s t-tests. We set the threshold for statistical significance at p□<□0.05.

## Results

### Cerebral Oxygenation Dynamics Under Awake and Isoflurane Conditions

During awake imaging, mice were securely head-fixed on an airflow-suspended Mobile HomeCage platform (Neurotar) that permitted spontaneous locomotion while ensuring stable PA image acquisition (Figure 1 C-E). To assess the usefulness of high-resolution ultrasound-photoacoustic (PA) imaging, we mapped the trajectory of cerebral oxygen saturation (sO_2_) across three distinct brain regions under awake conditions and during exposure to 2% isoflurane anesthesia (Figure 2A). Interestingly, sO_2_ levels were consistent across the three regions in the awake state (Figure 2B). Under isoflurane, however, we observed a significant increase in sO_2_ in Zone 1 (Figure 2C) and Zone 2 (Figure 2D), while Zone 3 remained relatively stable (Figure 2E). Notably, HbT levels, as quantified by PA imaging, remained unchanged throughout the experiment (Figure 2F–H).

**Figure 2.**
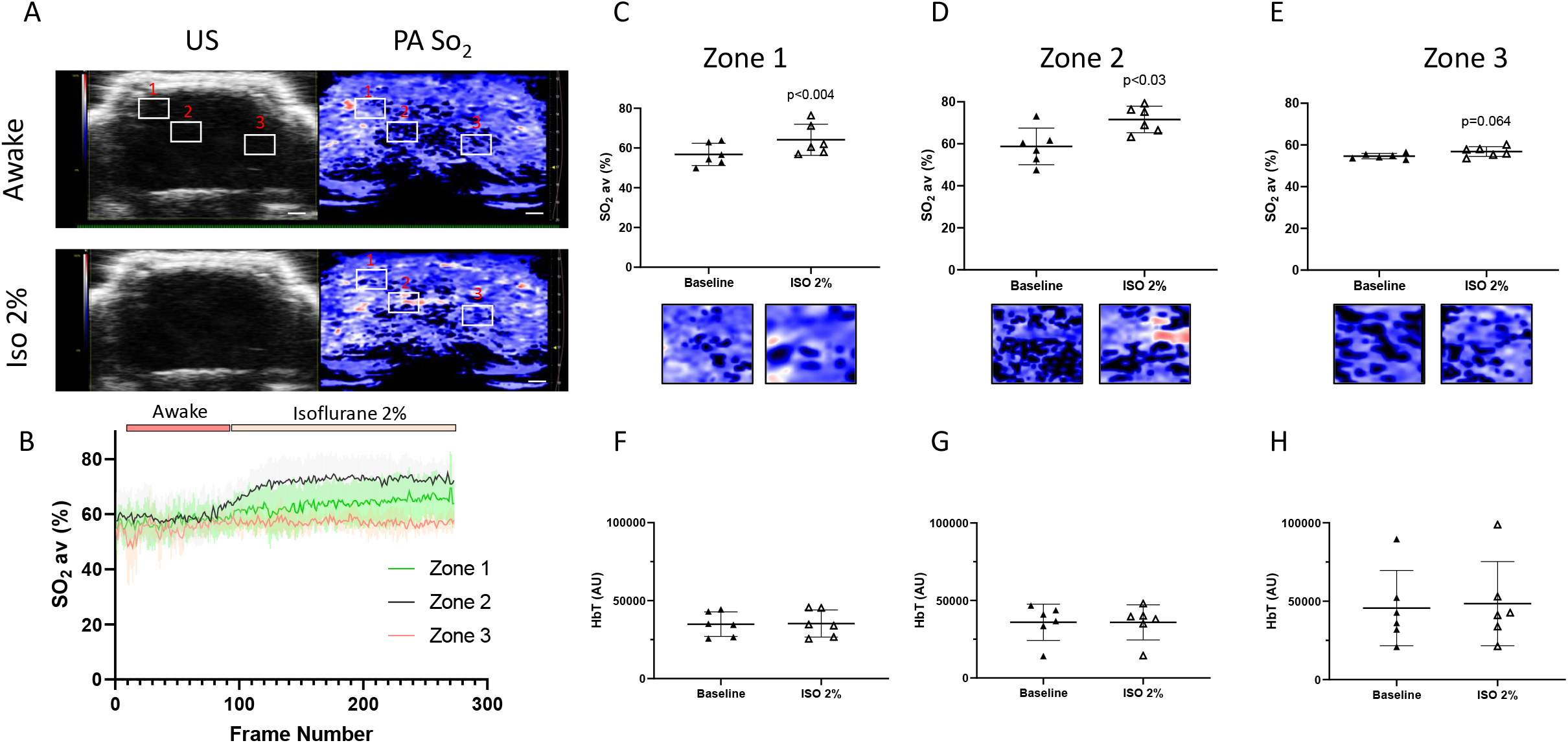
Photoacoustic brain imaging in awake versus anesthetized mice. **Panel A** Representative ultrasound (US) and photoacoustic (PA) images acquired in awake mice and under 2% isoflurane anesthesia. **Panel B** Cerebrovascular oxygen saturation (sO□) dynamics during awake and anesthetized states. **Panel C-E** Quantification of cerebrovascular sO□ in three distinct brain regions under both conditions. **Panel F-H** Quantification of total hemoglobin (HbT) in the same brain regions, highlighting anesthesia-dependent variations.

### Sensory Stimulation Evokes Localized Hemodynamic Responses in Awake Mice

We next investigated the changes in total hemoglobin (Hb) induced by a sensory stimulation task in awake, head-fixed mice. The right whiskers of mice positioned in the Mobile HomeCage were mechanically stimulated for 20–30 s periods while simultaneously acquiring PA signals in oxy-hemo Mode (Figure 3 A-B). Whisker stimulation elicited an increase in HbT within the contralateral (left) somatosensory barrel field cortex (n = 5, Figure 3C-D). The PA signal intensity increased rapidly following stimulus onset, reached a peak within a few seconds, and returned to baseline levels shortly after stimulus cessation, consistent with stimulus-evoked neurovascular coupling (Figure 3C). No significant changes in HbT were observed in ipsilateral regions or unrelated cortical areas (Figure 3D), confirming the regional specificity of the hemodynamic response.

**Figure 3.**
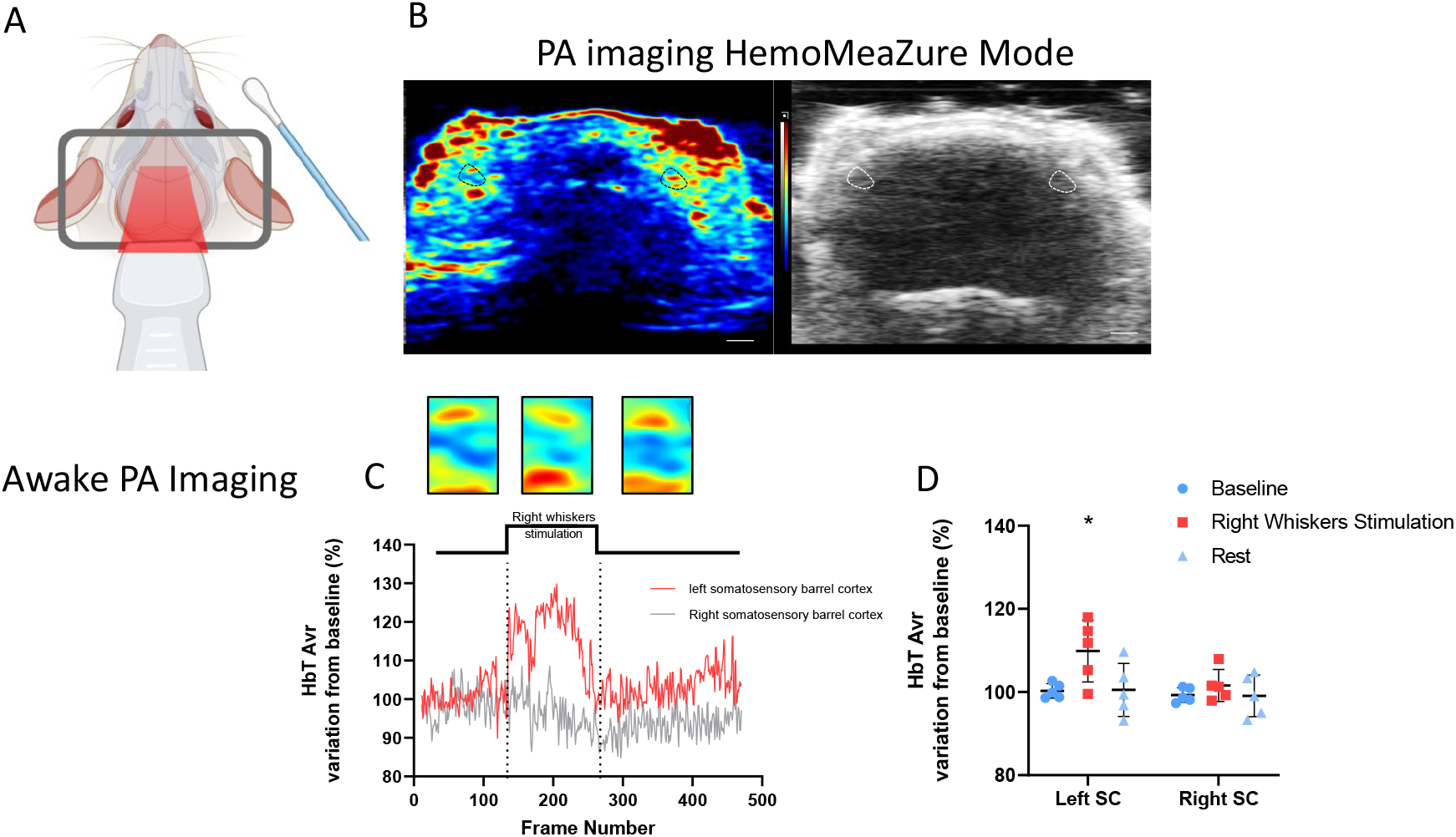
Photoacoustic imaging during right whisker stimulation in awake mice. Panel A Schematic of the whisker stimulation protocol and PA imaging setup. Panel B US and PA images used to monitor total hemoglobin trajectories during whisker stimulation. **Panel C** Temporal dynamics of cerebrovascular total hemoglobin (HbT) before, during, and after stimulation. **Panel D** Quantification of HbT changes across the same time windows, revealing hemodynamic responses to sensory input.

## Discussion

In this study, we demonstrate the feasibility of high-resolution ultrasound-photoacoustic imaging for non-invasive monitoring of cerebrovascular oxygenation dynamics in awake mice. By combining a customized head-fixation system with the Mobile HomeCage platform, we achieved stable acquisitions without requiring anesthesia or invasive cranial windows, thus minimizing confounding effects on neurovascular physiology.

Our findings confirm that exposure to 2% isoflurane induces a transient increase in cerebral sO_2_ across several brain regions, consistent with previous reports of isoflurane-induced cerebral vasodilation and increased oxygen delivery (11–13). Importantly, HbT levels remained stable, suggesting that the observed sO_2_ changes primarily reflect modifications in oxygen consumption rather than large-scale blood volume fluctuations. Furthermore, right whisker stimulation elicited robust, region-specific increases in HbT within the contralateral somatosensory barrel cortex, in agreement with well-established patterns of neurovascular coupling (14). These results validate the sensitivity of PA imaging to detect localized, stimulus-evoked hemodynamic responses in awake mice.

Compared with conventional techniques such as BOLD fMRI and fUS, PA imaging offers several advantages: it is non-invasive, cost-effective, and capable of directly quantifying oxy- and deoxyhemoglobin concentrations without the need for exogenous contrast agents. Unlike PA microscopy, which often requires cranial windows, our approach enables deep-brain imaging through the intact skull, making it suitable for longitudinal studies and translational neuroscience applications. Moreover, we observed that HbT signal changes measured with PA imaging were comparable to power Doppler responses measured with fUS, further validating our findings (15).

### Limitations

Despite its strengths, our study highlights several technical limitations of PA imaging. First, image resolution and penetration depth remain constrained compared with other modalities. Although our setup provided sufficient sensitivity to resolve cortical and subcortical structures, deeper brain regions remain challenging to access. Recent advances in tissue transparency techniques (16) may eventually extend the depth range of PA imaging for longitudinal studies specifically for cerebrovascular study.

Second, photoacoustic artifacts were occasionally observed in the reconstructed images (17), which can compromise data quality. Motion artifacts caused by spontaneous locomotion or subtle head muscle contractions during whisker stimulation resulted in transient image defocusing (18) and in our work could elevated PA noise. Adjusting acquisition parameters, such as increasing image persistence, can mitigate these effects, and light sedation has also been proposed to reduce spontaneous movements without abolishing neurovascular responses (1). Third, improving data quality and analysis speed remains an active area of development. Emerging deep learning-based reconstruction and denoising methods (19,20) show promise for reducing artifacts, enhancing spatial resolution, and accelerating quantitative PA data processing. Finally, while our study focused on the somatosensory barrel field cortex, a major tactile processing area activated by whisker stimulation in rodents (14), future work should explore PA imaging across distributed cortical and subcortical circuits to better capture network-level oxygenation dynamics.

## Conclusions

Overall, our results establish high-resolution ultrasound-photoacoustic imaging as a powerful, non-invasive tool for characterizing cerebrovascular oxygenation dynamics in awake mice. By enabling the detection of both baseline and stimulation-evoked hemodynamic responses without anesthesia or craniotomy, this approach opens new avenues for longitudinal studies of cerebral metabolism, sensory processing, and disease mechanisms. Integrating PA imaging with advanced motion correction, deep learning-based analysis, and emerging optical clearing strategies holds strong potential to further enhance its utility in preclinical neuroscience and translational research.

## Data Availability Statement

The data that support the findings of this study are available from the corresponding author upon reasonable request.

## Author Contributions

**Juri Aparicio Arias:** Investigation, Visualization, review & editing. **Chrystel Lafont:** Investigation, Methodology, Supervision, review & editing. **Philippe Trochet:** Conceptualization, Methodology, Writing – review & editing. **Dieter Fuchs:** Conceptualization, Methodology, Writing – review & editing. **Pierre Sicard:** Conceptualization, Investigation, Methodology, Supervision, Writing – original draft, review & editing.

## Acknowledgements

We gratefully thank the staff for animal housing (PhyMedExp). We acknowledge Imagerie du Petit Animal de Montpellier (IPAM) for accessing Photoacoustic high-resolution ultrasound (LRQA Iso9001; France Life Imaging (grant ANR-11-INBS-0006); IBISA; Leducq Foundation (RETP), I-Site Muse).

